# FastProtein – An automated software for *in silico* proteomic analysis

**DOI:** 10.1101/2023.12.19.572382

**Authors:** Renato Simões Moreira, Vilmar Benetti Filho, Guilherme Augusto Maia, Tatiany Aparecida Teixeira Soratto, Eric Kazuo Kawagoe, Bruna Caroline Russi, Luiz Claudio Miletti, Glauber Wagner

**Author notes:** Corresponding author: Laboratório de Bioinformática, Departamento de Microbiologia, Imunologia e Parasitologia, Centro de Ciências Biológicas, Universidade Federal de Santa Catarina, Campus João David Ferreira Lima. Setor F, Bloco G, Sala G809. Trindade, Florianópolis, SC, 88040-970 Brazil.

## Abstract

**Background:** Although various tools provide proteomic information, each has its limitations regarding execution platforms, libraries, versions, and data output format. Therefore, integrating data analyses generated using different software programs is a manual process that can prolong the analysis time.

**Results:** This paper presents FastProtein, a protein analysis pipeline tool developed in Java. This tool is user-friendly, easily installable, and provides important information regarding the subcellular location, transmembrane domains, signal peptide, molecular weight, isoelectric point, hydropathy, aromaticity, gene ontology, endoplasmic reticulum retention domains, and N-glycosylation domains of a protein. Furthermore, it helps determine the presence of glycosylphosphatidylinositol and obtain annotation information using InterProScan, PANTHER, PFam, and alignment-based annotation searches. Additionally, the software outputs a protein dataset with evidence of membrane localization.

**Conclusions:** The proposed tool provides the scientific community with an easy and user-friendly computational tool for proteomics data analysis. The tool is applicable to both small datasets and proteome-wide studies. It can be used in either the command line interface mode or through a web interface installed on a local server or via the BioLib web interface (http://biolib.com/UFSC/FastProtein). FastProtein also accelerates proteomics analysis routines by generating multiple results in a one-step run. The software is open-source and freely available. Installation and execution instructions, as well as the source code and test files generated for tool validation, are provided at https://github.com/bioinformatics-ufsc/FastProtein.

## 1. Background

Proteomic analysis generates a considerable amount of computational data that require bioinformatics analysis (Vaudel et al., 2016). Downstream analyses are required to assess the qualitative and quantitative features of proteins, which typically involve using several bioinformatics software packages individually and in a non-integrated manner (Chen et al., 2020; Jiménez-Munguía et al., 2018). Blast2GO (Conesa et al., 2005, p. 2) is an alternative tool widely used for functional annotation; however, its source code is closed, and users must buy a license to use it.

In contrast, FastProtein is publicly available and integrates functional annotations, database similarity searches, and protein feature predictions to enable global proteomic profiling in an automated and user-friendly manner. Furthermore, the detailed information obtained through FastProtein features can be used to search for proteins of interest.

## 2. Methods

### 2.1. Workflow

FastProtein uses protein FASTA file as input to generate protein profiles. The workflow analysis begins by cleaning the headers in the FASTA file to standardize the inputs for different software. The first step of FastProtein involves biochemical feature prediction, which provides attributes such as protein length, molecular mass (kDa), hydropathy, isoelectric point (PI), and aromaticity (Lobry & Gautier, 1994). These processes are executed using BioJava libraries (Lafita et al., 2019). Subsequently, FastProtein identifies the N-glycosylation and endoplasmic reticulum retention domains using the PROSITE (Sigrist et al., 2013) database entries PS00001 and PS00014, respectively.

Thereafter, the WoLF PSORT tool (Horton et al., 2007) is used to predict the subcellular locations of eukaryotic organisms. Subsequently, transmembrane site, signal peptide, and glycosylphosphatidylinositol (GPI)-anchored predictions are performed using the THMM-2 (Käll et al., 2007), SignalP 5.0, (Almagro Armenteros et al., 2019), and PredGPI (Pierleoni et al., 2008) tools, respectively. Additionally, the transmembrane domain and signal peptide are predicted using Phobius (Käll et al., 2004).

Functional annotations are optional and performed using InterProScan (v5.61-93.0) (Jones et al., 2014). The outputs are merged and parsed to obtain the PFam (Mistry et al., 2021) and PANTHER (Thomas et al., 2022) domains, InterPro (IPR) annotations (Jones et al., 2014), and gene ontology (GO) terms (Ashburner et al., 2000). The sets of GO terms associated with the protein are determined by analyzing all databases using InterProScan. These terms are organized into a file that can be imported into the WEGO 2.0 (Ye et al., 2018) platform for complementary analysis. This platform is used to group, visualize, compare, and generate GO plots.

Ontology terms provide quantitative reports on molecular functions, cellular components, and biological processes. This process generates a file containing the GO terms (one line per protein, followed by GO terms in table-separated files).

Another important step in this workflow is similarity analysis. Using the FASTA file provided by the user as a database, FastProtein returns the best hit for each protein (with its identity and coverage percentage) using the BLASTp (Camacho et al., 2009) or DIAMOND (Buchfink et al., 2021) algorithms.

FastProtein provides six pieces of evidence for membrane protein identification: GPI-anchored, two predictions for transmembrane domains (TM, predicted via TMHMM-2.0; and PHOBIUS_TM, predicted via Phobius), subcellular localization predicted via WoLF PSORT (SL), GO, and IPR annotations. The last two pieces are based on two text files that can be customized by the user. In this case, each protein receives a value from 0–6 and a set of evidence. Finally, a report file and a FASTA file are generated for the proteins with membrane evidence.

### 2.2. FastProtein Output Files

FastProtein outputs quantitative and qualitative files, as elucidated in **Supplementary Table 1**. Additionally, it generates an integrated histogram and scatter plot of molecular masses and isoelectric points, as well as a bar chart that displays the predicted subcellular localizations. The images are created at 300 DPI using Matplotlib (Hunter, 2007) and Seaborn (Waskom, 2021) libraries.

Individual protein information is provided in tab-separated values (TSV), comma-separated values (CSV), text-formatted (TXT), XLS (Microsoft Excel), open document format (ODF), and separated image (SEP) file formats. SEP is a custom format similar to that output by ProtComp v9 (http://www.softberry.com/).

Every generated file is stored in a temporary directory located within the FastProtein installation directory named using a universally unique identifier (UUID) created at the beginning of the run, which enables parallel execution. At the end of the process, the temporary directory is renamed as the directory selected by the user (with a default value of fastprotein _results). In case of processing errors, the previously generated files can be reused by employing the -cdt directory> command, which indicates the directory from which the generated files must be copied. This option is only available in the command line interface (CLI) mode. Additionally, users can use a command (-zip) to compress the results in a zip file, which deletes the output directory and only retains the Zip file. By default, the BioLib Web execution generates the Zip file comprising the results. If FastProtein is executed using the BioLib CLI, the resultant file is stored in the execution directory (BioLib_results). An execution log is saved in the output directory, and the logging level can be set in the CLI mode.

A web module was developed using Python (v3.9.2) and Flask (v2.2.3) to run FastProtein in a Docker container. This module receives a webpage request, executes FastProtein, and allows downloading the generated result in Zip format from the same page. Starting the Docker container results in the automatic launching of the module at port 5000.

Finally, all third-party software are configured such that the user can execute them individually. Thus, the FastProtein Docker container becomes a bioinformatics suite with several pre-installed software packages (Table 1) and is ready to be executed even without FastProtein.

**Table 1.** Software used in the FastProtein pipeline.

| **Software Library** | **Purpose** | **Reference** |
| --- | --- | --- |
| WoLF PSORT | Subcellular location (for eukaryotes only) | doi:10.1093/nar/gkm259 |
| TMHMM-2 | Transmembrane predictions of domain sites | doi:10.1093/nar/gkm256 |
| SignalP 5.0 | Signal peptide prediction and location of cleavage sites in protein | doi:10.1038/s41587-019-0036-z |
| InterProScan | Functional predictions (Ontology terms) | doi:10.1093/nar/gkac240 |
| BioJava | Bioinformatic support library | doi:10.1371/journal.pcbi.1006791 |
| Docker | Lightweight linux containers | doi/10.5555/2600239.2600241 |
| Phobius | A combined transmembrane topology and signal peptide predictor | 10.1016/j.jmb.2004.03.016 |
| PredGPI | GPI-Anchor Predictor | 10.1186/1471-2105-9-392 |
| BLASTp | Sequence aligner for proteins | 10.1186/1471-2105-10-421 |
| DIAMOND | Sequence aligner for proteins | 10.1038/s41592-021-01101-x |
| Seaborn | Python data visualization library | 10.21105/joss.03021 |
| Matplotlib | Python library for creating static, animated, and interactive visualizations | 10.1109/MCSE.2007.55 |

Fig. 1(a) shows the FastProtein workflow, which indicates the software execution order and the corresponding outputs generated.

**Fig. 1.**
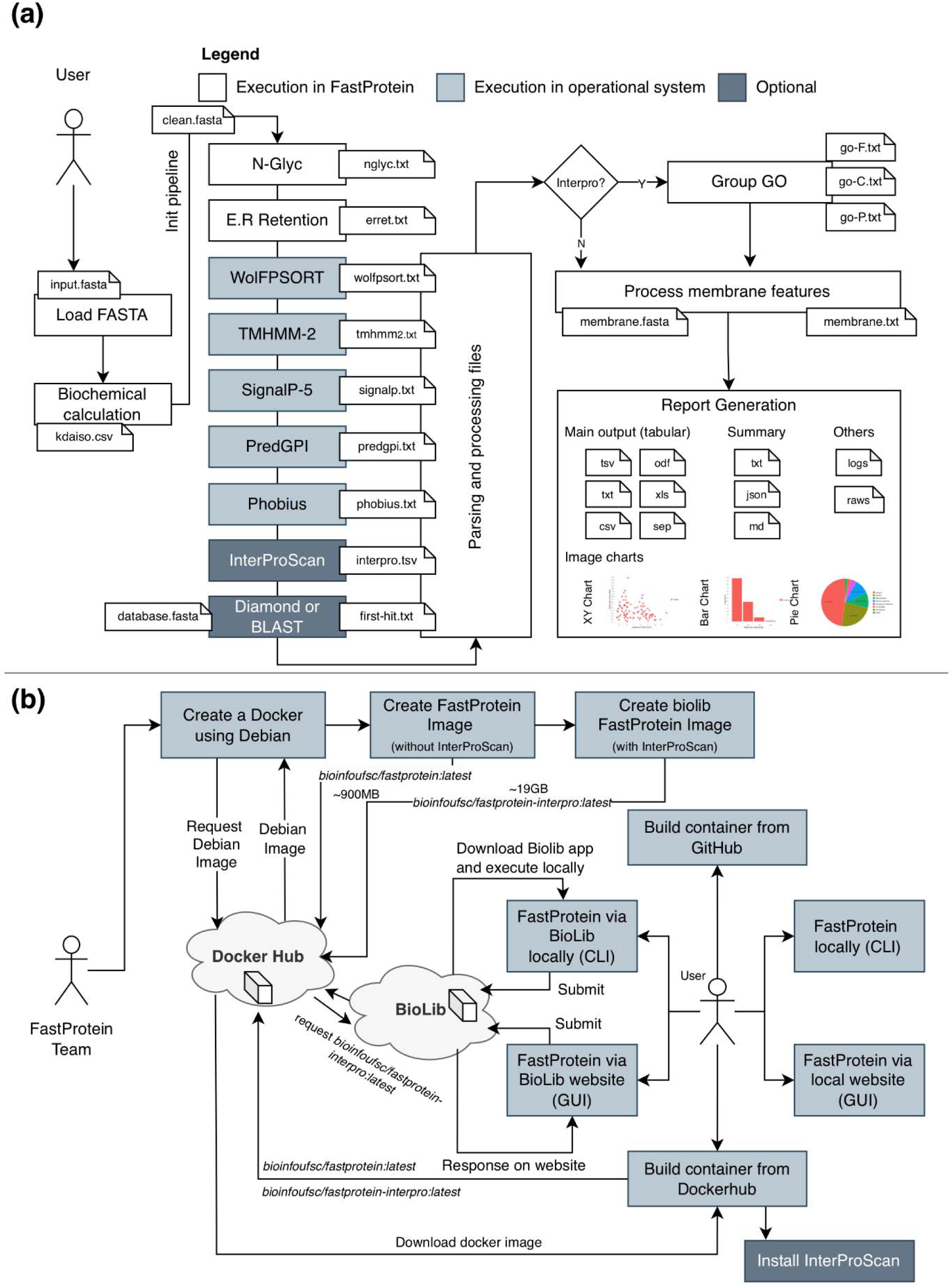
(a) FastProtein pipeline: starting with the user-provided FASTA file, the proteins are processed and sent to third-party software. Thereafter, the results are converted and merged, GO terms are counted, proteins with membrane evidence are evaluated, and graphs and files are generated. (b) Computational structure of FastProtein: the Docker with Debian Linux is customized using third-party software, and two versions of FastProtein files are sent to the DockerHub (with and without InterProScan). Subsequently, the Docker with InterProScan is associated with the BioLib project, providing the user with several implementation options.

### 2.3. Computational Infrastructure

A Debian-based Docker image (Merkel, 2014) is available at https://hub.docker.com/r/bioinfoufsc/fastprotein. This image is 900 MB (compressed) and includes an installation script for InterProScan (v5.61-93.0), which is required for functional annotation (recommended). Another custom image (used on the BioLib platform) was created based on the first image, but with InterProScan already installed. This image had a total size of approximately 18 GB (compressed); it can be downloaded from https://hub.docker.com/r/bioinfoufsc/fastprotein-interpro. The third-party software and dependencies employed are listed in **Table 1**. The computational organization of this infrastructure is illustrated in Figure 1(b).

FastProtein was developed using the Java 17 programming language and is available at BioLib (https://BioLib.com). The installation guide, usage instructions, and the source code are available at https://github.com/bioinformatics-ufsc/FastProtein. FastProtein can be executed in four different ways: (1) through the web-based Graphical User Interface (GUI) in the same Docker container, which involves building an image using a Dockerfile or downloading an already prepared image (Fig. 2(a)); (2) through the CLI from a local Docker container; (3) through the CLI, which requires both Python (3.7 or higher) and the BioLib package (pyBioLib) to be installed; and (4) through the BioLib Web GUI (Fig. 2(b)), available at https://BioLib.com/UFSC/FastProtein.

**Fig. 2.**
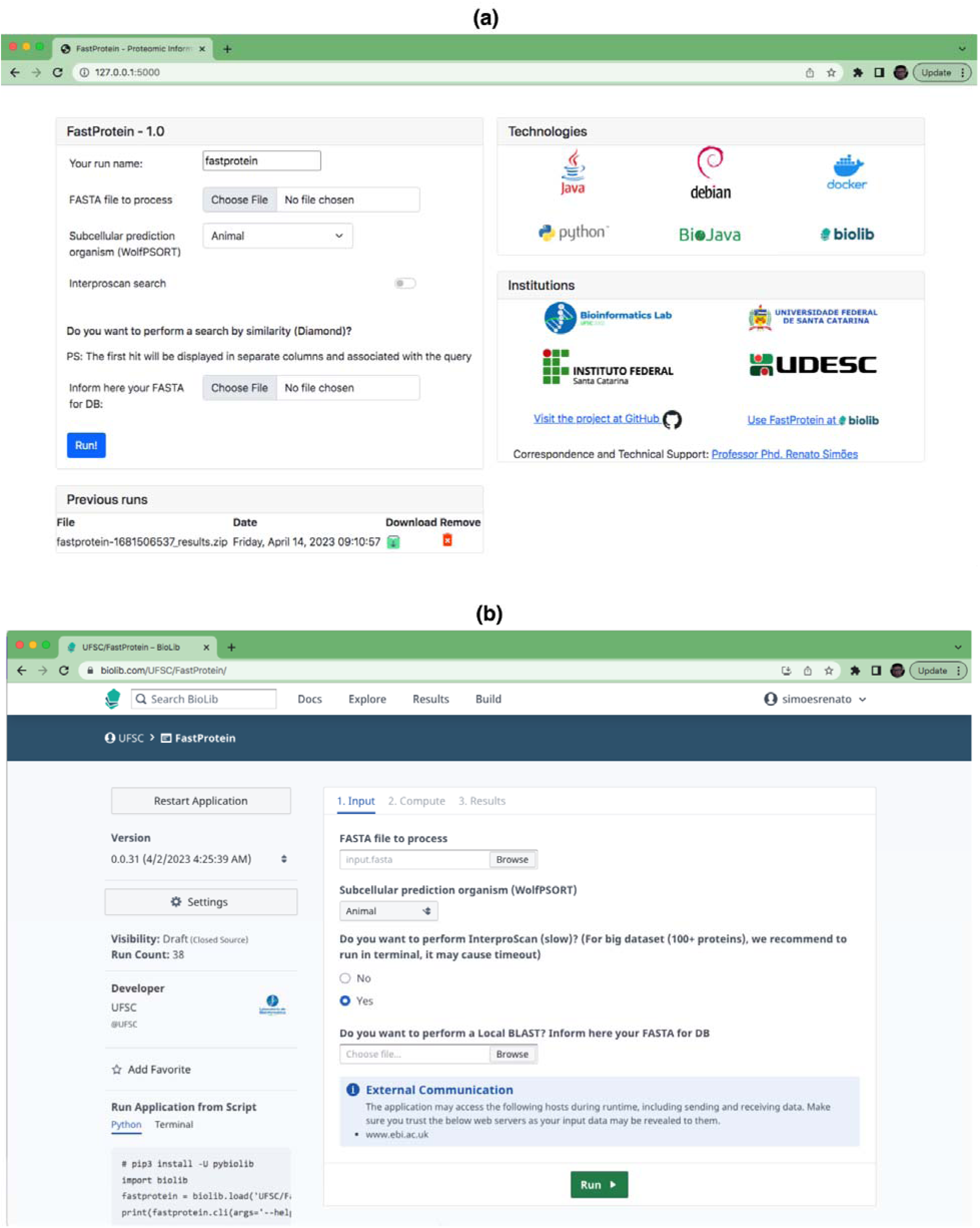
(a) Main GUI of the FastProtein web module deployed on a local server. The embedded web server is started at http://<ip_server>:5000, which can be accessed using any web browser. (b) BioLib Web interface after implementation displaying information and results of the run, and a link for downloading the zipped file is made available automatically.

**Fig. 3.**
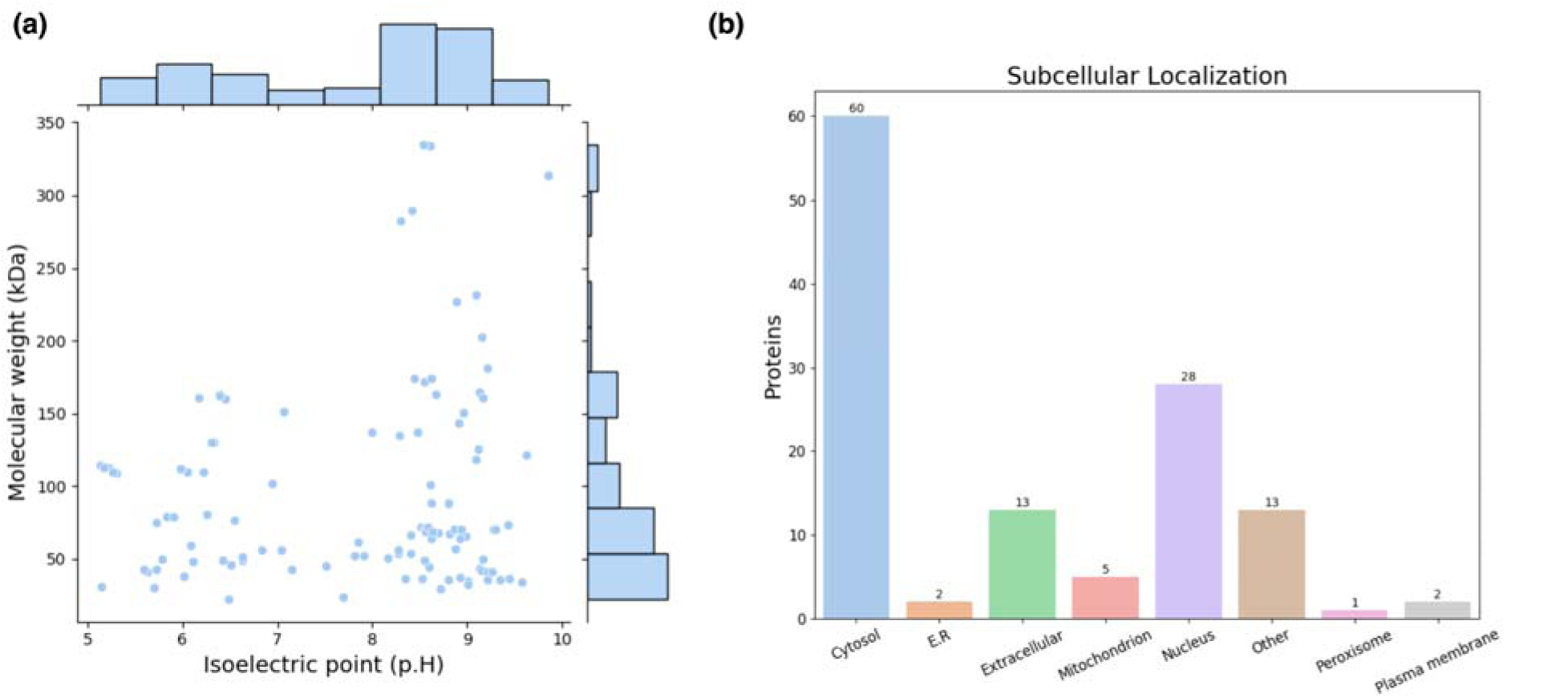
(a) Histogram illustrating the molecular weight and isoelectric point on a scatter plot. (b) Bar graph of subcellular locations. All graphics include 124 of 125 proteins of *P. malariae* (on of them was rejected) tested on FastProtein and generated using Seaborn (Waskom, 2021) and Matplotlib (Hunter, 2007) libraries.

BioLib Web has an input limitation for proteins, which can result in a timeout of the platform itself. Therefore, it is recommended that CLI or the web version of FastProtein should be used for analyses involving more than 200 proteins.

### 2.4. Experiments

To demonstrate the FastProtein workflow, analyses were performed on a subset of 125 proteins from the *Plasmodium malariae* proteome (UP000219813). This test run was performed in the CLI mode using DIAMOND and BLASTp for local alignment.

For proteome-wide analysis and performance benchmarking, we used the following proteomes (strain, Proteome ID) downloaded on April 2, 2023, from UniProt (The UniProt Consortium et al., 2021): *Plasmodium vivax* (strain Salvador I, UP000008333), *Trypanosoma brucei* (strain 927/4 GUTat10.1, UP000008524), *Cryptosporidium muris* (strain RN66, UP000001460), *Toxoplasma gondii* (strain ATCC 50861 / VEG, UP000002226), *Aspergillus novofumigatus* (strain IBT 16806, UP000234474), and *Cyanidioschyzon merolae* (strain NIES- 3377 / 10D, UP000007014). All proteomes were analyzed using the DIAMOND and BLASTp alignment methods (against the respective proteomes), and each proteome was run in triplicate. The WoLF PSORT dataset was set up to consider the closest organism, as the WoLF PSORT models are restricted to animals, plants, and fungi.

## 3. Results

From the first dataset comprising 125 proteins, only 124 were deemed eligible for analysis, with protein A0A1A8WBL6 excluded due to the presence of “X” in its sequence. The total analysis time for this dataset, using DIAMOND as the local aligner (default parameters), was 2 minutes and 58 seconds, with 5 seconds exclusively required by FastProtein and the remaining time by third-party software. When BLASTp was used as the local aligner, the total execution time was 7 minutes and 13 seconds, with 3 minutes and 49 seconds used by BLASTp for computations, 5 seconds by FastProtein, and the remaining by other third-party software.

The average molecular weight and isoelectric point in the *Plasmodium malariae* dataset were 98.29 ± 71.30 kDa and 7.79 ± 1.36, respectively. The distribution generated by FastProtein is shown in Fig. 2(a). The average hydrophilicity and aromaticity were -0.50 ± 0.31 and 0.09 ± 0.03, respectively. Out of the analyzed proteins, 30 were predicted to contain transmembrane domains, three exhibited GPI anchoring, and an additional 30 were estimated to be membrane proteins. Furthermore, among the latter group, four exhibited multiple pieces of evidence supporting their localization to the membrane (GO, IPR, PHOBIUS_TM, and TM). Among the 124 *P. malariae* proteins, 11 were predicted to have signal peptides and 36 had ER retention domains, with the KEEL and KNEL domains being the most frequently occurring, existing in seven proteins each. Of the 124 proteins, 123 had N-glycosylation domains, with the NNS domain occurring most frequently (116 proteins). The subcellular locations are shown in Fig. 2(b), wherein Cytosol and Nucleus are the most frequent ones, with 60 and 28 proteins, respectively.

Considering proteome-wide rounds, the fastest analysis was performed on the *C. muris* dataset (3,930 proteins, with 3,924 processed and 6 ignored) using the DIAMOND alignment method with an execution time of 28.85 ± 5.74 minutes. The slowest analysis was performed on the *T. gondii* dataset (8,404 proteins) using the BLASTp alignment method, with an execution time of 262.35 ± 2.63 minutes. By changing the aligner from BLASTp to DIAMOND, the execution time was decreased to 77.75 ± 8.90 minutes. Interestingly, even though *A. novofumigatus* had 3,076 more proteins than *T. gondii*, it took 97.42 ± 0.21 minutes to analyze it using the BLASTp method and 67.71 ± 1.73 minutes using the DIAMOND method. The best result in terms of proteins analyzed per minute was obtained for the *A. novofumigatus* dataset with 170 proteins (using DIAMOND), whereas the worst was obtained for *T. gondii* with 32 proteins (using BLASTp). In all DIAMOND executions, the analysis was completed within a few seconds, unlike in BLASTp, where the fastest execution required 8.02 ± 0.05 minutes and the slowest required 193.06 ± 0.92 minutes to complete. Only the execution times of the results obtained with different aligners differed, and considerably similar results were obtained using different algorithms (Hernández-Salmerón & Moreno-Hagelsieb, 2020).

The average execution times for each software used in the pipeline were as follows: WoLF PSORT (∼2 minutes), TMHMM-2.0 (∼8 minutes), SignalP5 (∼2 minutes), PredGPI (∼2 minutes), Phobius (∼11 minutes), InterProScan (∼25 minutes), DIAMOND (less than 1 minute), and BLASTp (∼52 minutes). The average time required for the internal execution of FastProtein, file generation, conversions, and calculations was approximately 1 minute. Both files generated from the subset of *P. malariae* proteins and proteome-wide rounds are available at https://github.com/bioinformatics-ufsc/FastProtein/ (including the intermediate files, which were removed during processing). The remaining data are presented in <u>Supplementary Material 1</u>.

## Discussion

FastProtein is a user-friendly and easy-to-install protein analysis pipeline tool that provides important information regarding protein datasets. FastProtein integrates calculations of molecular weight, isoelectric point, hydropathy, and aromaticity with predictions of subcellular location, transmembrane domains, signal peptide and GPI-anchor, GO, endoplasmic reticulum retention, and N-glycosylation domains, as well as analyses using InterProScan, PANTHER, PFam, and alignment-based annotation searches. Additionally, the software provides a dataset of proteins with evidence of membrane localization, which is important for immunogenicity studies during vaccine development (Cheng et al., 2021; Kis et al., 2018) and diagnostic tools, such as ELISA (de Haro-Cruz et al., 2019; Iha et al., 2022) and western blotting (Begum et al., 2022; Crescitelli et al., 2021; Springhorn & Hoppe, 2019).

FastProtein outputs files in formats that are widely used in the scientific community, including TSV, XLS, and ODF, as well as high-quality 300 DPI images, which is a widely used standard that does not require additional parsing.

The total execution time of FastProtein depends on the InterProScan analysis and the optional alignment method selected. By default, DIAMOND was selected owing to its relatively faster execution time, although BLASTp is also available. The global median time for proteome-wide analyses was approximately 116 proteins per minute. By using the DIAMOND alignment method, this was increased to approximately 142 proteins per minute, whereas the BLASTp method yielded approximately 90 proteins per minute. Thus, proteomic data from a large dataset can be quickly obtained using FastProtein.

The only requirement for using the FastProtein software is the installation of Docker or BioLib for local runs. By default, executions through BioLib Web have no limitations; however, unlike the CLI version, the platform has a timeout restriction that can interrupt prolonged processing. Thus, FastProtein is a viable tool for researchers with no background in bioinformatics analysis because it provides a user-friendly interface similar to well-established software such as Blast2GO (Conesa et al., 2005), MEGA11 (Tamura et al., 2021), and MaxQuant (Prianichnikov et al., 2020). Additionally, it contributes to the initiatives that aim to democratize access to bioinformatics, such as the BioLib (http://biolib.com) and Galaxy (Sheynkman et al., 2014) projects.

## Conclusions

FastProtein is a novel and user-friendly pipeline tool for proteomic data analysis. It is applicable to both small datasets and proteome-wide studies. Furthermore, it can be used either in the CLI mode or through a web interface installed on a local server or that available on BioLib (http://biolib.com/UFSC/FastProtein). FastProtein accelerates proteomics analysis routines by generating multiple results in a one-step run. The software is open-source and available at https://github.com/bioinformatics-ufsc/FastProtein, along with installation and execution instructions and test files generated for validation.

## Availability and requirements

**Project name:** FastProtein

**Project home page:** https://github.com/bioinformatics-ufsc/FastProtein

**Operating system(s):** Platform independent

**Programming language:** Java 17+

**Other requirements:** Docker or BioLib

**License:** Apache 2.0

**Any restrictions to use by non-academics:** No additional restrictions.

## Abbreviations

CSV: Comma-separated values
DPI: Dots per inch
GB: Gigabytes
GO: Gene ontology
GPI: Glycosylphosphatidylinositol
GUI: Graphical user interface
IPR: InterProScan
MB: Megabytes
ODF: Open document format
SEP: Custom format where each line represents a protein information
SL: Subcellular localization
TM: Transmembrane domain
TSV: Tab-separated values
TXT: text-formatted
UUID: Universally unique identifier
XLS: Microsoft Excel

## Acknowledgements

FastProtein was developed and tested at the Laboratório de Bioinformática, located in the Departamento de Microbiologia, Imunologia e Parasitologia (MIP), Universidade Federal de Santa Catarina (UFSC), in partnership with the Instituto Federal de Santa Catarina (IFSC) and Universidade do Estado de Santa Catarina (UDESC), Brazil. The authors declare that this study was conducted in the absence of any commercial or financial relationships that could be construed as potential conflicts of interest.

## Author information

**Instituto Federal de Santa Catarina (IFSC), Campus Gaspar, Gaspar, Brazil.**

Renato Simões Moreira, Bruna Caroline Russi

**Laboratório de Bioinformática, Universidade Federal de Santa Catarina (UFSC), Florianópolis, Brazil.**

Renato Simões Moreira, Vilmar Benetti Filho, Guilherme Augusto Maia, Tatiany Aparecida Teixeira Soratto, Eric Kazuo Kawagoe, Bruna Caroline Russi, Glauber Wagner

Laboratório de Hemoparasitas e Vetores, Centro de Cie□ncias Agroveterinárias **(CAV), Universidade do Estado de Santa Catarina (UDESC), Lages, Brazil.**

Luiz Claudio Miletti

## Contributions

This work was conceived and initially implemented by RSM. The organizational aspects, computational infrastructure, control versions, and script implementations were performed by VBF and EKK. BCR developed the web module. GAM and TATS were responsible for testing, whereas GW, LCM, and RSM analyzed the results and wrote the manuscript. All the authors have read and approved the final manuscript.

## Corresponding author

Glauber Wagner

## Ethics declarations

### Ethics approval and consent to participate

Not applicable.

### Consent for publication

Not applicable

## Supplementary information

**Supplementary Table 1.** Summary of files output by the FastProtein pipeline.

| File | Purpose |
| --- | --- |
| output.[xls, odf, csv, tsv, txt, sep] | Individual information of each protein |
| output.md | Summary in Markdown containing totalization, charts, GO, subcellular localization, and the first ten proteins |
| summary.txt | Summary containing information such as totalization, subcellular localization, ER retentions domain, and N-glycosylation domains |
| summary.json | File containing the totalizations and all individual protein information |
| raw/membranes.txt | File containing proteins with at least one membrane evidence, and indication of the type of evidence |
| raw/membrane.fasta | FASTA file containing proteins with at least one membrane evidence |
| raw/ignored.fasta | FASTA file containing ignored proteins |
| go-[C, F, P].txt | File containing GO totalization by GO-Term |
| console.log | Run output |
| wego.txt | File in native WEGO format, it can be loaded at https://wego.genomics.cn/ |
| image/ | Directory containing a histogram of molecular mass and isoelectric point integrated with a scatter plot and bar chart of subcellular localization predicted via WoLF PSORT (300 DPI available) |
| raw/ | Directory containing all raw data generated using the third-party software |

## References

Ashburner, M., Ball, C. A., Blake, J. A., Botstein, D., Butler, H., Cherry, J. M., Davis, A. P., Dolinski, K., Dwight, S. S., Eppig, J. T., Harris, M. A., Hill, D. P., Issel-Tarver, L., Kasarskis, A., Lewis, S., Matese, J. C., Richardson, J. E., Ringwald, M., Rubin, G. M., & Sherlock, G. (2000). Gene Ontology: Tool for the unification of biology. Nature Genetics, 25(1), 25–29. 10.1038/75556

Begum, H., Murugesan, P., & Tangutur, A. D. (2022). Western blotting: A powerful staple in scientific and biomedical research. BioTechniques, 73(1), 58–69. 10.2144/btn-2022-0003

Buchfink, B., Reuter, K., & Drost, H.-G. (2021). Sensitive protein alignments at tree-of-life scale using DIAMOND. Nature Methods, 18(4), 366–368. 10.1038/s41592-021-01101-x

Camacho, C., Coulouris, G., Avagyan, V., Ma, N., Papadopoulos, J., Bealer, K., & Madden, T. L. (2009). BLAST+: Architecture and applications. BMC Bioinformatics, 10(1), 421. 10.1186/1471-2105-10-421

Chen, X.-L., Liu, C., Tang, B., Ren, Z., Wang, G.-L., & Liu, W. (2020). Quantitative proteomics analysis reveals important roles of N-glycosylation on ER quality control system for development and pathogenesis in *Magnaporthe oryzae*. PLoS Pathogens, 16(2), e1008355. 10.1371/journal.ppat.1008355

Cheng, K., Zhao, R., Li, Y., Qi, Y., Wang, Y., Zhang, Y., Qin, H., Qin, Y., Chen, L., Li, C., Liang, J., Li, Y., Xu, J., Han, X., Anderson, G. J., Shi, J., Ren, L., Zhao, X., & Nie, G. (2021). Bioengineered bacteria-derived outer membrane vesicles as a versatile antigen display platform for tumor vaccination via Plug-and-Display technology. Nature Communications, 12(1), 2041. 10.1038/s41467-021-22308-8

Conesa, A., Götz, S., García-Gómez, J. M., Terol, J., Talón, M., & Robles, M. (2005). Blast2GO: A universal tool for annotation, visualization and analysis in functional genomics research. *Bioinformatics (Oxford*, England), 21(18), 3674–3676. 10.1093/bioinformatics/bti610

Crescitelli, R., Lässer, C., & Lötvall, J. (2021). Isolation and characterization of extracellular vesicle subpopulations from tissues. Nature Protocols, 16(3), 1548–1580. 10.1038/s41596-020-00466-1

de Haro-Cruz, M. J., Guadarrama-Macedo, S. I., López-Hurtado, M., Escobedo-Guerra, M. R., & Guerra-Infante, F. M. (2019). Obtaining an ELISA test based on a recombinant protein of Chlamydia trachomatis. International Microbiology, 22(4), 471–478. 10.1007/s10123-019-00074-4

Hernández-Salmerón, J. E., & Moreno-Hagelsieb, G. (2020). Progress in quickly finding orthologs as reciprocal best hits: Comparing blast, last, diamond and MMseqs2. BMC Genomics, 21(1), 741. 10.1186/s12864-020-07132-6

Hunter, J. D. (2007). Matplotlib: A 2D Graphics Environment. Computing in Science & Engineering, 9(03), 90–95. 10.1109/MCSE.2007.55

Iha, K., Tsurusawa, N., Tsai, H.-Y., Lin, M.-W., Sonoda, H., Watabe, S., Yoshimura, T., & Ito, E. (2022). Ultrasensitive ELISA detection of proteins in separated lumen and membrane fractions of cancer cell exosomes. Analytical Biochemistry, 654, 114831. 10.1016/j.ab.2022.114831

Jiménez-Munguía, I., Pulzova, L., Kanova, E., Tomeckova, Z., Majerova, P., Bhide, K., Comor, L., Sirochmanova, I., Kovac, A., & Bhide, M. (2018). Proteomic and bioinformatic pipeline to screen the ligands of *S. pneumoniae* interacting with human brain microvascular endothelial cells. Scientific Reports, 8(1), 5231. 10.1038/s41598-018-23485-1

Jones, P., Binns, D., Chang, H.-Y., Fraser, M., Li, W., McAnulla, C., McWilliam, H., Maslen, J., Mitchell, A., Nuka, G., Pesseat, S., Quinn, A. F., Sangrador-Vegas, A., Scheremetjew, M., Yong, S.-Y., Lopez, R., & Hunter, S. (2014). InterProScan 5: Genome-scale protein function classification. Bioinformatics, 30(9), 1236–1240. 10.1093/bioinformatics/btu031

Käll, L., Krogh, A., & Sonnhammer, E. L. L. (2004). A Combined Transmembrane Topology and Signal Peptide Prediction Method. Journal of Molecular Biology, 338(5), 1027– 1036. 10.1016/j.jmb.2004.03.016

Kis, Z., Shattock, R., Shah, N., & Kontoravdi, C. (2018). Emerging Technologies for LowlJCost, Rapid Vaccine Manufacture. Biotechnology Journal, 1800376. 10.1002/biot.201800376

Lobry, J. R., & Gautier, C. (1994). Hydrophobicity, expressivity and aromaticity are the major trends of amino-acid usage in 999 *Escherichia coli* chromosome-encoded genes. Nucleic Acids Research, 22(15), 3174–3180. 10.1093/nar/22.15.3174

Merkel, D. (2014). Docker: Lightweight linux containers for consistent development and deployment. Linux Journal, 2014(239), 2.

Mistry, J., Chuguransky, S., Williams, L., Qureshi, M., Salazar, G. A., Sonnhammer, E. L. L., Tosatto, S. C. E., Paladin, L., Raj, S., Richardson, L. J., Finn, R. D., & Bateman, A. (2021). Pfam: The protein families database in 2021. Nucleic Acids Research, 49(D1), D412–D419. 10.1093/nar/gkaa913

Prianichnikov, N., Koch, H., Koch, S., Lubeck, M., Heilig, R., Brehmer, S., Fischer, R., & Cox, J. (2020). MaxQuant Software for Ion Mobility Enhanced Shotgun Proteomics. Molecular & Cellular Proteomics, 19(6), 1058–1069. 10.1074/mcp.TIR119.001720

Sheynkman, G. M., Johnson, J. E., Jagtap, P. D., Shortreed, M. R., Onsongo, G., Frey, B. L., Griffin, T. J., & Smith, L. M. (2014). Using Galaxy-P to leverage RNA-Seq for the discovery of novel protein variations. BMC Genomics, 15(1), 703. 10.1186/1471-2164-15-703

Sigrist, C. J. A., de Castro, E., Cerutti, L., Cuche, B. A., Hulo, N., Bridge, A., Bougueleret, L., & Xenarios, I. (2013). New and continuing developments at PROSITE. Nucleic Acids Research, 41(Database issue), D344–347. 10.1093/nar/gks1067

Springhorn, A., & Hoppe, T. (2019). Western blot analysis of the autophagosomal membrane protein LGG-1/LC3 in Caenorhabditis elegans. In Methods in Enzymology (Vol. 619, pp. 319–336). Elsevier. 10.1016/bs.mie.2018.12.034

Tamura, K., Stecher, G., & Kumar, S. (2021). MEGA11: Molecular Evolutionary Genetics Analysis Version 11. Molecular Biology and Evolution, 38(7), 3022–3027. 10.1093/molbev/msab120

The UniProt Consortium, Bateman, A., Martin, M.-J., Orchard, S., Magrane, M., Agivetova, R., Ahmad, S., Alpi, E., Bowler-Barnett, E. H., Britto, R., Bursteinas, B., Bye-A-Jee, H., Coetzee, R., Cukura, A., Da Silva, A., Denny, P., Dogan, T., Ebenezer, T., Fan, J., … Teodoro, D. (2021). UniProt: The universal protein knowledgebase in 2021. Nucleic Acids Research, 49(D1), D480–D489. 10.1093/nar/gkaa1100

Thomas, P. D., Ebert, D., Muruganujan, A., Mushayahama, T., Albou, L.-P., & Mi, H. (2022). PANTHER: Making genome-scale phylogenetics accessible to all. Protein Science: A Publication of the Protein Society, 31(1), 8–22. 10.1002/pro.4218

Vaudel, M., Verheggen, K., Csordas, A., Ræder, H., Berven, F. S., Martens, L., Vizcaíno, J. A., & Barsnes, H. (2016). Exploring the potential of public proteomics data. PROTEOMICS, 16(2), 214–225. 10.1002/pmic.201500295

Waskom, M. (2021). seaborn: Statistical data visualization. Journal of Open Source Software, 6(60), 3021. 10.21105/joss.03021

Ye, J., Zhang, Y., Cui, H., Liu, J., Wu, Y., Cheng, Y., Xu, H., Huang, X., Li, S., Zhou, A., Zhang, X., Bolund, L., Chen, Q., Wang, J., Yang, H., Fang, L., & Shi, C. (2018). WEGO 2.0: A web tool for analyzing and plotting GO annotations, 2018 update. Nucleic Acids Research, 46(W1), W71–W75. 10.1093/nar/gky400

